# Phosphate starvation regulates cellulose synthesis to modify root growth

**DOI:** 10.1101/2023.09.15.557998

**Authors:** Ghazanfar Abbas Khan, Arka Dutta, Allison Van de Meene, Kristian EH. Frandsen, Michael Ogden, James Whelan, Staffan Persson

## Abstract

In the model plant *Arabidopsis thaliana*, the absence of the essential macro-nutrient phosphate reduces primary root growth through decreased cell division and elongation, requiring alterations to the polysaccharide-rich cell wall surrounding the cells. Despite its significance, the regulation of cell wall synthesis in response to low phosphate levels is not well understood. In this study, we show that plants increase cellulose synthesis in roots under limiting phosphate conditions, which leads to changes in the thickness and structure of the cell wall. These changes contribute to the reduced growth of primary roots in low phosphate conditions. Furthermore, we found that the cellulose synthase complex activity at the plasma membrane increases during phosphate deficiency. Moreover, we show that this increase in the activity of the cellulose synthase complex is likely due to alterations in the phosphorylation status of cellulose synthases in low phosphate conditions. Specifically, phosphorylation of CELLULOSE SYNTHASE 1 at the S688 site decreases in low phosphate conditions. Phosphomimic versions of CELLULOSE SYNTHASE 1 with an S688E mutation showed significantly reduced cellulose induction and primary root length changes in low phosphate conditions. Protein structure modelling suggests that the phosphorylation status of S688 in CELLULOSE SYNTHASE 1 could play a role in stabilizing and activating the cellulose synthase complex. This mechanistic understanding of root growth regulation under limiting phosphate conditions provides potential strategies for changing root responses to soil phosphate content.

## Introduction

Phosphorus, an essential nutrient in all life forms, can only be obtained by plants as inorganic phosphate (Pi). In many environments, both native and managed, the amount of Pi in the soil is below that required to support plant growth; thus, plants have developed various mechanisms to increase their survival during Pi limitation. These mechanisms include the production of carboxylic acids to release complexed Pi, expressing high-affinity transporters to maximize Pi uptake, and altering metabolism to prioritize essential processes (Chiou & Lin, 2011). Moreover, the root system architecture (RSA) of *Arabidopsis thaliana* undergoes significant changes in response to low Pi levels (Abel, 2017). When Pi is limited, the plant promotes the growth of secondary roots and root hairs while halting the growth of primary roots. This adaptation allows the plant to increase its root volume in the topsoil, where Pi is more readily available, in an effort to maximize the acquisition of this essential nutrient (Paz-Ares *et al*., 2022). Thus, RSA remodelling results in a morphological adaption to enable survival under Pi starvation conditions. The recent argument that root growth inhibition in low Pi conditions is not a biological response but rather a result of artificial root exposure to blue light that triggers photo-Fenton chemistry has been challenged by (Gao *et al*., 2021). They showed that blue light illumination in shoots leads to reduced primary root growth under Pi-deficient conditions through long-distance shoot-to-root signalling.

The transcription factor PHOSPHATE STARVATION RESPONSE 1 (PHR1) plays a major role in coordinating most of the plant’s metabolic responses to low internal Pi (Bustos *et al*., 2010). By contrast, RSA remodelling is influenced by external Pi availability and is controlled by the gene LOW PI ROOT 1 (LPR1) and its homologue LPR2 (Abel, 2017). LPR1, a cell wall ferroxidase in the cell wall, is involved in Fe redox cycling and the production of reactive oxygen species (ROS) and callose in response to low Pi (Naumann *et al*., 2022b). These cell wall changes restrict cell expansion and division. Here, it is thought that ROS and cell wall modifications work together to stiffen the cell wall (Balzergue *et al*., 2017).

Plant cell walls are composed of various polysaccharides, including cellulose, hemicelluloses, callose, and pectins, and play a critical role in directing cell growth and shaping the plant (Lampugnani *et al*., 2018). Every plant cell forms a primary cell wall, which dynamically expands during growth, but once cellular growth ceases, specific cells, such as tracheary elements and fibres, deposit thick secondary cell walls to support the cell against stress (Nicolas *et al*., 2022). Cellulose, consisting of β-(1→4)-D-glucan chains, forms the framework of the cell wall and is synthesized by cellulose synthase complexes (CSCs) in the plasma membrane(Polko & Kieber, 2019).

In the Arabidopsis genome, there are 10 *CESA* genes, with *CESA1*, *CESA3*, *CESA6*, and *CESA2*, *CESA5*, and *CESA9* involved in primary cell wall formation, and *CESA4*, *CESA7*, and *CESA8* involved in secondary cell wall formation (Richmond, 2000, Persson *et al*., 2007, Taylor *et al*., 2003). The function of *CESA10* is suggested to be redundant to *CESA1* in the seed coat (Griffiths *et al*., 2015). The Arabidopsis CESAs have a 60% sequence identity and contain seven transmembrane helices, with the catalytic domain of the CESAs located between the second and third transmembrane helices (Purushotham *et al*., 2020). This domain contains a canonical motif and is surrounded by two plant-specific domains, the plant-conserved region (PCR) and class-specific region (CSR), which play important roles in CESA oligomerization and CSC assembly (Atanassov *et al*., 2009). Multiple oligomeric states of CESAs, including homodimers, have been reported and it is hypothesized that their oligomerization plays a role in the early stages of cellulose CSC assembly (Atanassov et al., 2009). CSCs move at a speed of 200-350 nm/min on the plasma membrane and are responsible for synthesizing and depositing cellulose microfibrils (Paredez *et al*., 2006). The movement of CSCs is thought to be driven by the catalytic activity of the complexes and is regulated by various factors, including protein phosphorylation of the CESA proteins (Chen *et al*., 2010, Sánchez-Rodríguez *et al*., 2017). While sub-cellular processes that underpin the guidance of cellulose biosynthesis to establish cell shape and organs are increasingly better understood (Lampugnani et al., 2018), a mechanistic connection between the perception of low nutrient abundance, such as Pi deficiency, and a defined modulation of cell walls, required for changes in RSA, is not yet known.

Here, we show that low Pi levels lead to increased cellulose deposition in roots. These changes correspond to a decrease in phosphorylation of CESA1 at the S688 site in the CSR domain, which likely regulates CESA1 activity to mediate root growth and cellulose deposition during Pi starvation. Our findings suggest that the regulation of cellulose synthesis plays a role in regulating root growth in response to low Pi.

## Results

### Cellulose synthesis is induced in plant roots in response to low Pi

To understand cell wall changes in response to low Pi, we quantified cellulose in the roots of seedlings grown on Pi deficient (20 μM) and Pi sufficient (1 mM) conditions. The roots grown on low Pi accumulate significantly higher levels of cellulose compared to the ones grown on sufficient Pi, suggesting that cellulose synthesis is influenced by Pi availability (Fig. 1A). Because we observed an increase in cellulose deposition, we next assessed whether the cell wall thickness was also changed in response to Pi starvation. We, therefore, prepared root sections for transmission electron microscopy and imaged cell walls of cortex cells in the root elongation zone. We found that the corresponding cell walls in Pi-deficient conditions were significantly thicker than those grown in sufficient Pi conditions (Fig. 1BC). To understand the role of cellulose changes in root growth regulation, we next measured the primary root length of the mildly cellulose deficient mutant *procuste1-1* (*prc1-1*) (Fagard *et al*., 2000) under Pi deficient conditions. As *prc1-1* mutants already have shorter roots as compared to Col-0 when grown on Pi sufficient media, it is difficult to make a direct comparison for changes in primary root length in response to Pi deficiency. Therefore, we calculated the ratio of primary root length of seedlings grown on Pi deficient media as compared to Pi sufficient media for each genotype. The *prc1-1* mutant roots are constitutively shorter than those of the wild type, and they do not show a significant change in their root growth when exposed to low Pi conditions. (Fig. 1DEF). These findings suggest that controlling cellulose synthesis is likely necessary for altering root growth in response to low Pi conditions.

**Figure 1.**
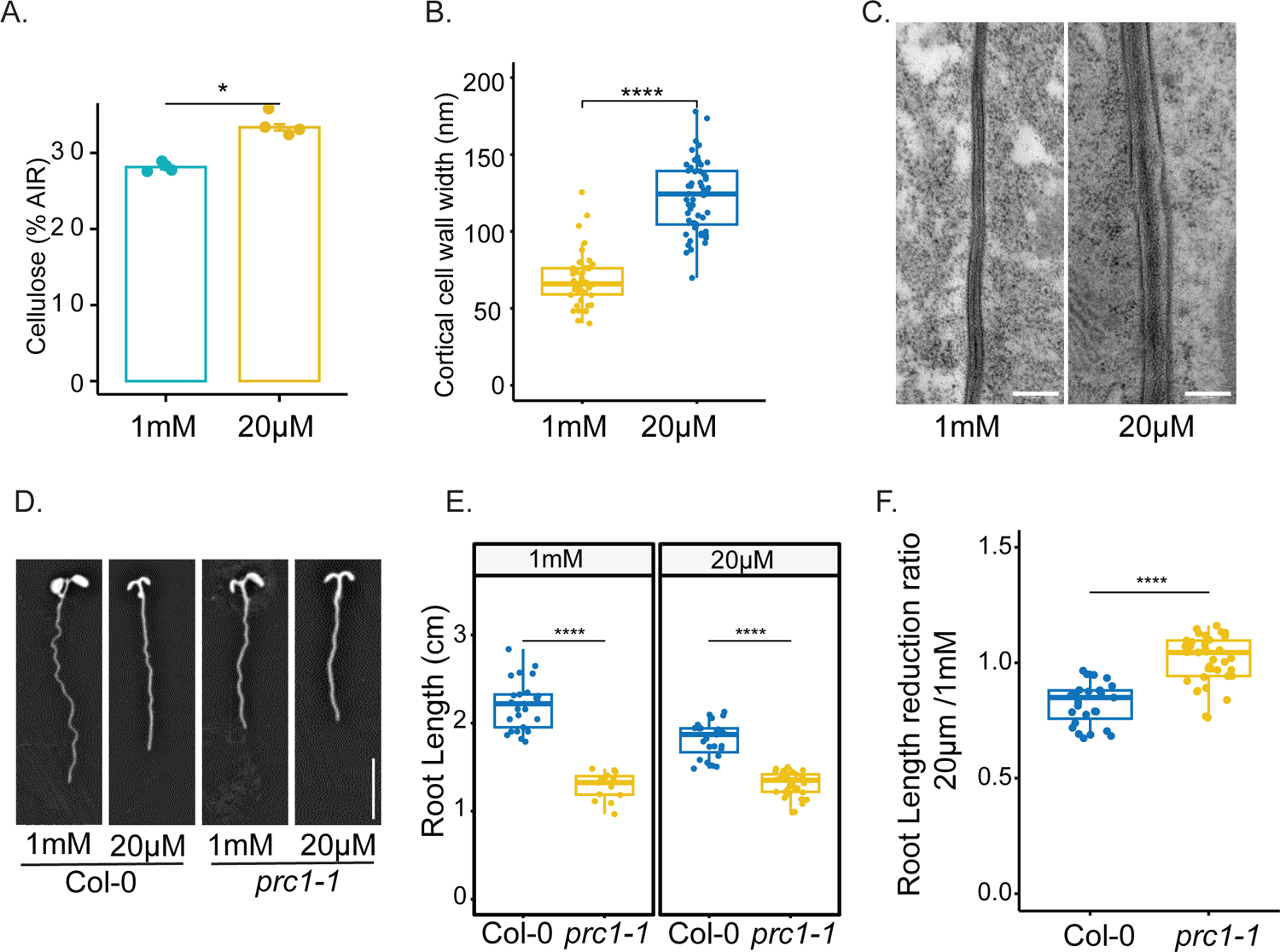
Cellulose deposition is increased in roots grown under Pi deficient conditions. **A.** Cellulose quantification of 8-day-old roots grown on Pi sufficient (1mM) or Pi deficient (20µM) conditions. Data is presented as means ± SE, n =4. AIR: Alcohol insoluble residue **B.** The thickness of the cell walls in cortical cells within the root elongation zone was quantified using transmission electron microscopy (TEM). This analysis was conducted on plants that had been grown for 8 days under either Pi-sufficient or Pi-deficient conditions. Six seedlings from each of these conditions were used for the quantification of cell wall thickness. **C.** Representative TEM images of cell walls between cortical cells. Scale bar = 200nm. **D.** Representative images of 8 days old Col-0 and *prc1-1* roots grown on Pi sufficient and Pi deficient conditions. Scale bar = 1cm **E.** Root length of 8 days old seedlings grown on Pi sufficient and Pi deficient media. **F.** Ratio of primary root length reduction of plants grown in Pi deficient conditions as compared to plants grown in Pi sufficient conditions. Asterisks denote statistical significance (*, P < 0.05; **, P < 0.01; ***, P < 0.001 and; ****, P < 0.0001) according to Student’s t test. The graphics and statistical analysis were generated using the ggpubR package in R.

### Primary wall CESA levels do not change under Pi deficiency

To gain further insights into cellulose regulation under Pi deficiency, we quantified the expression of primary wall *CESA*s in Arabidopsis roots grown under Pi sufficient and Pi deficient conditions. Primary wall *CESA* genes did not show any changes in their expression in response to Pi deficiency (Fig. 2A), indicating that post-transcriptional changes are likely responsible for the observed cellulose regulation. By contrast, the Pi transporter *PHOSPHATE TRANSPORTER 1;3* (*PHT1;3)* (Nussaume *et al*., 2011) showed a significant expression increase (Fig. 2A), confirming that roots were indeed experiencing a Pi deficiency. To determine if changes in CESA protein levels instead could contribute to the cellulose changes, we performed shotgun proteomics analyses and performed label-free quantification of proteins. We identified 7798 proteins from 44795 peptide groups. We found that 232 proteins were differentially regulated in response to Pi starvation. Proteins commonly associated with Pi starvation, such as PURPLE ACID PHOSPHATASE 17 (PAP17), NON-SPECIFIC PHOSPHOLIPASE C4 (NPC4), and PHT1;3 were induced, indicating that the roots were experiencing Pi starvation. In contrast, proteins involved in RNA translation were repressed (Table S1). The analysis of differentially abundant proteins revealed an enrichment of GO terms related to Pi ion homeostasis, protein phosphorylation and translation (Fig. 2C). The alteration in translation observed may be due to the reallocation of resources in low Pi conditions, which is in line with previous observations. (Cheng *et al*., 2021). However, we found no changes in the abundance of primary wall CESA proteins in response to Pi deficiency (Fig. 2B). These results suggest that primary wall CESA protein levels do not change in response to Pi deficiency.

**Figure 2:**
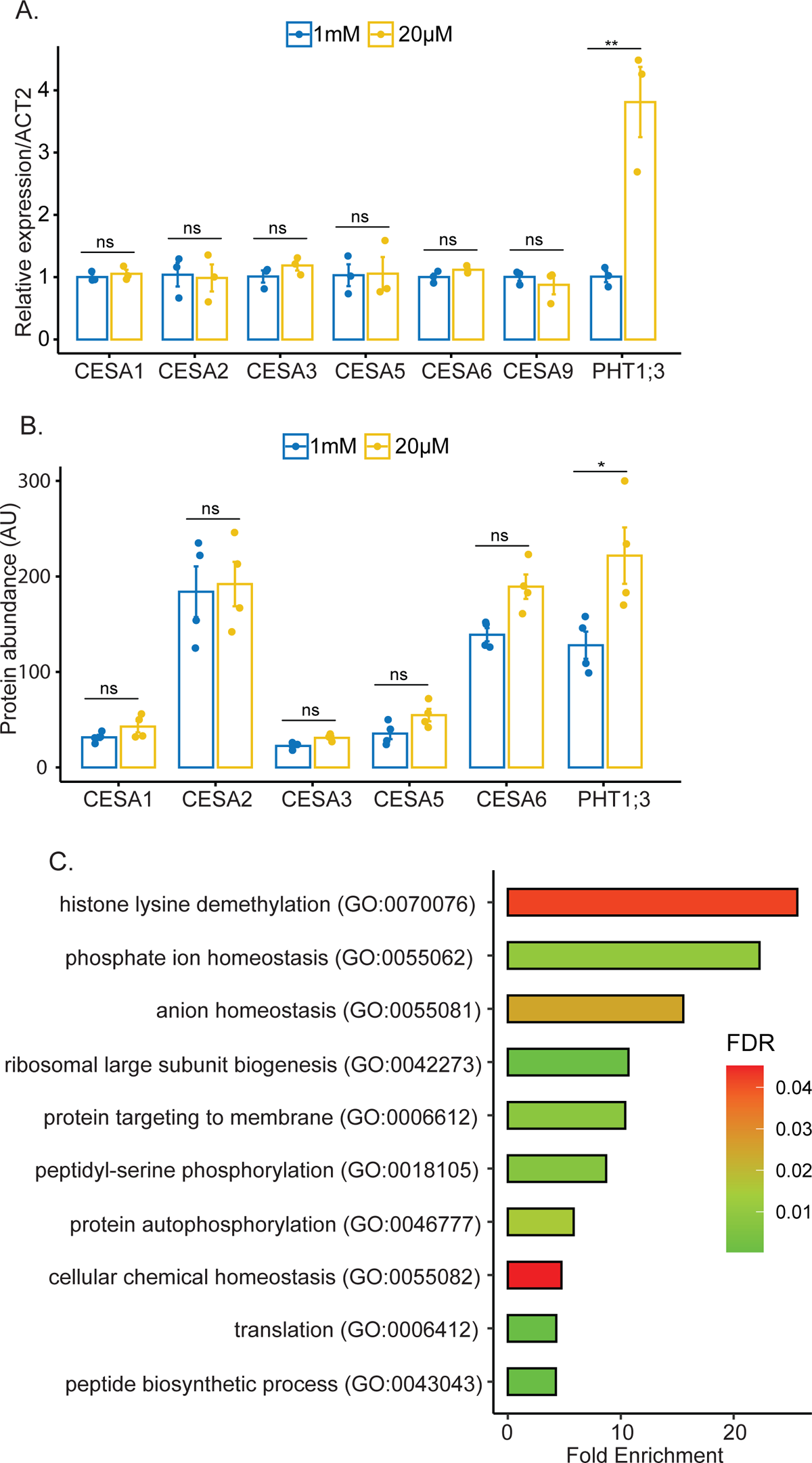
Primary wall CESA transcript and protein levels under Pi deficiency. **A.** Relative transcript abundance of primary wall *CESA*s and *PHT1;3* (Marker gene for Pi deficiency) in the eightday-old roots of Arabidopsis seedlings grown under Pi sufficient (1mM) and Pi deficient (20µM) conditions. Data are presented as means ± SE, n =3. **B.** Protein abundance of primary wall CESAs and PHT1;3 in the roots of Arabidopsis plants grown in Pi-sufficient and Pi-deficient conditions. Data are presented as means ± SE, n =4. **C.** Go term enrichment of differentially expressed proteins in Pi deficient media compared to Pi sufficient media. Asterisks denote statistical significance (ns, non-significant; *, P < 0.05; **, P < 0.01) according to Student’s t-test. The graphics and statistical analysis were generated using the ggpubR package in R.

### Cellulose synthase activity is increased in response to low Pi

To synthesize cellulose, CSCs move in the plasma membrane, and their speed is thought to be dependent on the catalytic activity of the CESAs (Pedersen *et al*., 2023). To test if the cellulose changes in response to Pi deficiency were due to the modulation of CESA activity, we estimated the CESA speed at the plasma membrane. Indeed, we found that the speed of CESAs was significantly faster in the elongating root cells of plants grown on low Pi media compared to plants grown on optimum Pi media (Fig. 3A, B). This indicates that the increase in cellulose deposition in low Pi-grown plants is at least partially due to increased CESA activity at the plasma membrane. The density of CSCs at the plasma membrane did not differ significantly between roots grown on Pi-deficient media and those grown on sufficient Pi media. (Fig. 3C). Overall, these results show that Pi availability regulates CESA activity in root cells.

**Figure 3:**
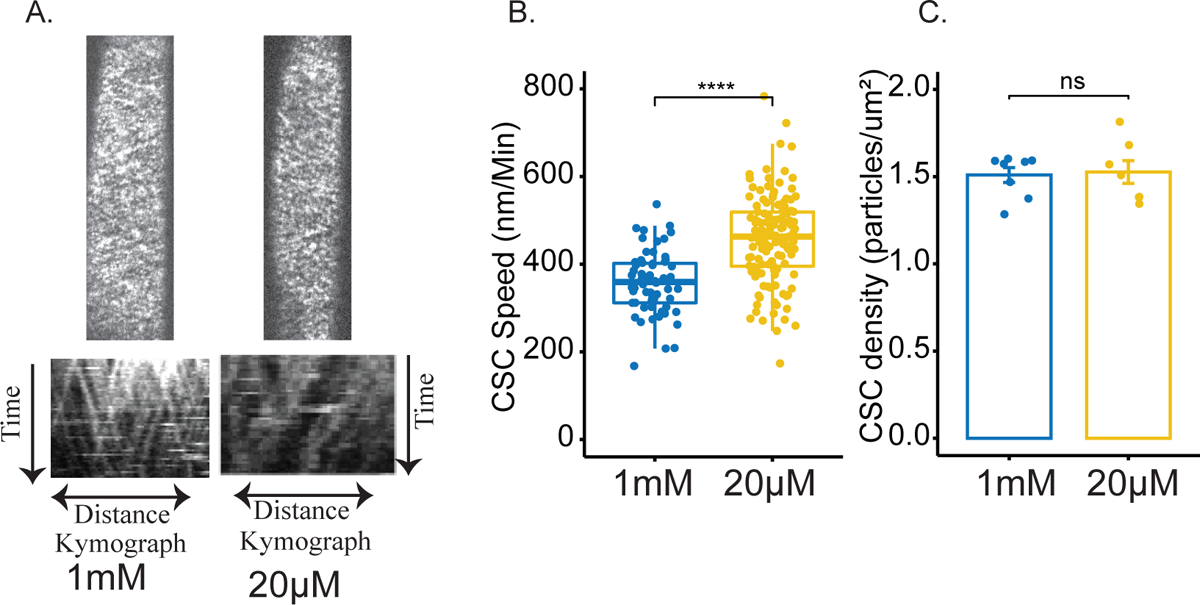
CSC activity increases under low Pi conditions. Roots were grown on agar plates containing Pi sufficient (1mM) or Pi deficient media (20µM). **A.** Plasma membrane localization and kymographs of YFP-CesA6 images taken from cells in the root elongation zone of eight day-old Arabidopsis seedlings. **B.** Boxplots showing CSC speeds in the elongation zone of roots of seedlings grown on Pi sufficient or deficient media. **C.** Bar plots showing CSC density at the plasma membrane of root cells in seedlings grown in Pi sufficient or deficient media. Data are presented as means ± SE, n >7. Asterisks denote statistical significance (ns, non-significant; *, P < 0.05; **, P < 0.01; ***, P < 0.001 and; ****, P < 0.0001) according to Student’s t test. The graphics and statistical analysis were generated using the ggpubR package in R.

### CESA1 is differentially phosphorylated under low Pi conditions

There is compelling evidence to suggest that CSC speed, and thus CESA activity, is affected by the phosphorylation status of the CESAs (Sanchez-Rodriguez *et al*., 2017, Speicher *et al*., 2018b). We examined the impact of low Pi on protein phosphorylation using shotgun proteomics and phospho-enriched peptide quantification. Our results revealed 7,982 phosphorylated peptides out of 44,795 total peptides, with 6,932 being serine phosphorylations, 1,016 being threonine phosphorylations, and 34 being tyrosine phosphorylations. Of these, 6,908 were single phosphorylation, 971 were double phosphorylation, and 103 were triple phosphorylation (Fig 4. AB, Table S2). A total of 552 peptides belonging to 422 proteins were differentially phosphorylated (Adj. p-value ≤0.05) in response to Pi starvation. The data showed an overrepresentation of GO terms associated with the auxin signaling, biosynthesis of phosphatidylinositol phosphate, miRNA processing, the detection of abiotic stimuli, and root development among differentially phosphorylated peptides in response to Pi starvation (Fig. 4C). Quantification of CESA phosphorylation showed significant changes in response to low Pi treatments as compared to sufficient Pi. Specifically, we observed a reduction in phosphorylation at the S11 and S688 sites in CESA1, as well as a decrease in the abundance of a triply phosphorylated peptide from CESA3 under these conditions (Table S2). We focused on the role of phosphorylation at the S688 site in CESA1, as it likely plays a role in CSC complex activity and stability (Fig. 6). Additionally, we were able to identify the corresponding unphosphorylated site, which showed an increase in abundance, further supporting the changes in phosphorylation at this site in response to Pi deficiency (Fig. 5AB). These results indicate that phosphorylation at S688 of CESA1 is regulated in response to Pi deficiency, indicating that CESA activity might be regulated through changes in phosphorylation status in response to low Pi.

**Figure 4:**
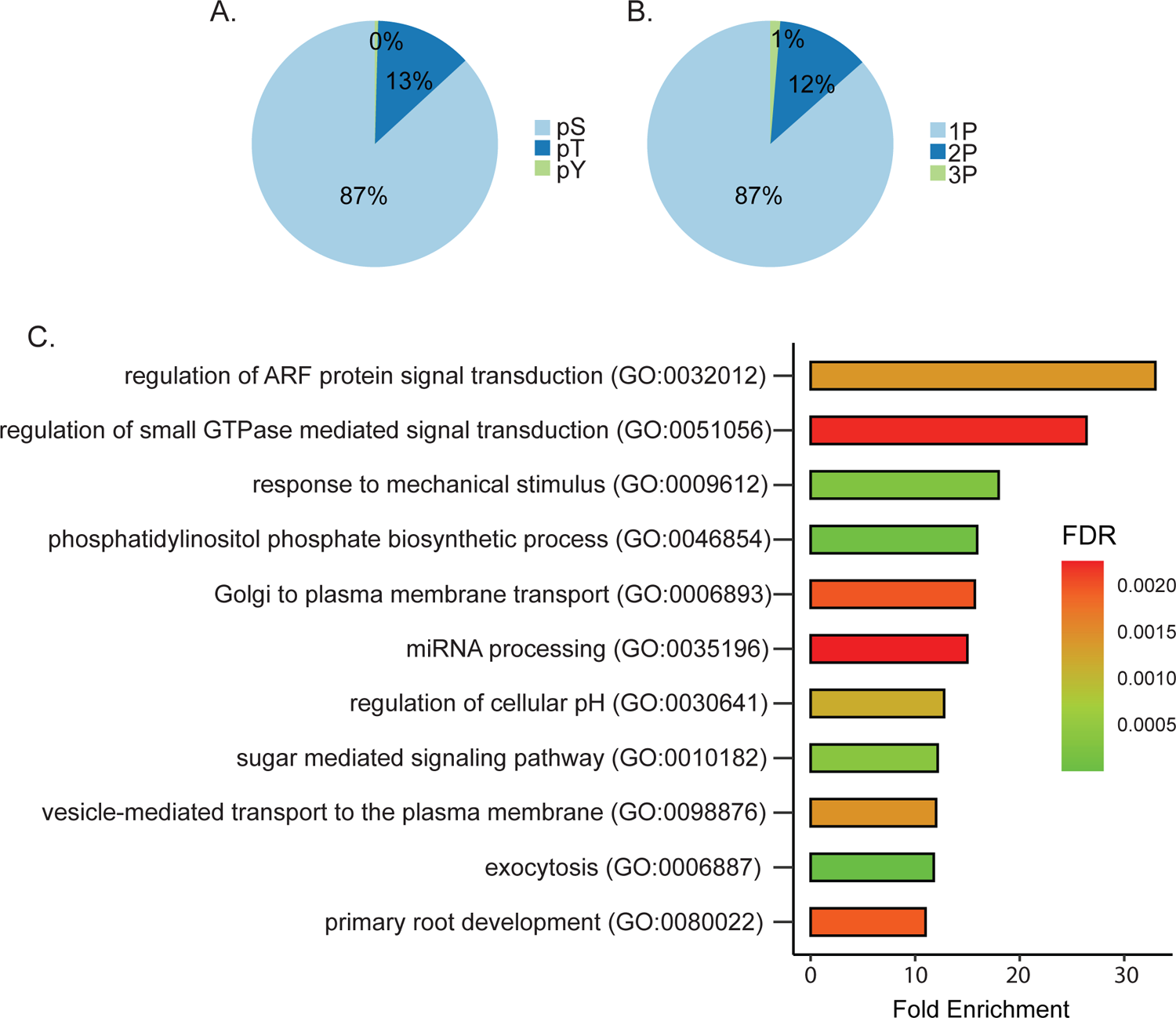
Changes in protein phosphorylation in response to Pi starvation. Plants were grown in a hydroponics system for 14 days with sufficient Pi (1mM) and then transferred to either Pi sufficient (1mM) or Pi deficient (20µM) media for 48 hours before experiments. **A.** Pie chart displaying the percentage of serine, threonine, and tyrosine phosphorylation in our data. **B.** Pie chart showing the percentage of single, double, and triply phosphorylated peptides in our data. **C.** GO term enrichment of differentially phosphorylated proteins in Pi deficient media in comparison to Pi sufficient media.

### Changes in CESA1 phosphorylation regulate root growth under Pi deficiency

To understand the role of the S688 phosphosite in Pi deficiency-mediated root growth regulation, we performed site-directed mutagenesis of S688 into phopsho-null (S688A) or phosphomimic (S688E) and transformed these versions, including wild type version (S688S) into *cesa1* knockout mutant. Plants with the wild-type version of CESA1 fully rescued the lethal phenotype of *cesa1* mutants (SAIL_278_E08) (Fig. 5C). We recovered three lines of heterozygous *cesa1* mutants containing the S688 phospho-null version of CESA1. These lines were allowed to self-pollinate, and 30 descendent plants from each of these lines were genotyped. However, we were unable to recover any homozygous *cesa1* mutant lines. These results suggest that S688 phospho-null version of CESA1 is unable to rescue the gametophytic lethal phenotype of *cesa1* mutants. By contrast, the phosphomimetic version was able to fully complement the *cesa1* mutant, indicating that some degree of phosphorylation at site S688 is critical for CESA1 function. The measurement of primary root length in plants segregating for both the *cesa1* mutation and phosphomimic T-DNA revealed no significant differences between Pi sufficient and Pi deficient growth conditions (Fig. S1). However, under Pi deficient conditions, we did observe some degree of segregation in terms of primary root length (Fig. S1). It is important to note that CESA1 forms functional cellulose synthase complexes in conjunction with other CESA proteins. In the presence of multiple forms of CESA proteins, determining which protein forms are incorporated into a functional cellulose synthase complex becomes challenging. To establish a more controlled experimental setting, we specifically isolated plants that carried the phospho-mimic version in a homozygous *cesa1* (-/-) background. Primary root length measurement of phosphomimetic plants in the homozygous *cesa1* (-/-) background grown under Pi deficient conditions showed significantly reduced changes in primary root length compared to the wild type (Fig. 5CDE). Moreover, S688E plants did not show an increase in cellulose in their roots when grown on Pi deficient conditions (Fig. 5F). These results indicate that phosphorylation at phosphosite S688 likely regulates CESA1 activity in response to Pi deficiency. To understand the impact of S688 phosphorylation on CSC activity, we crossed the phosphomimetic plants with YFP-CESA6 to enable CSC imaging in this background. However, these plants experienced silencing, likely due to the presence of three T-DNAs (SAIL_278_E08, S688E and YFP-CESA6), and we were unable to recover phosphomimetic plants with a bright enough YFP-CESA6 marker suitable for CSC imaging.

**Figure 5:**
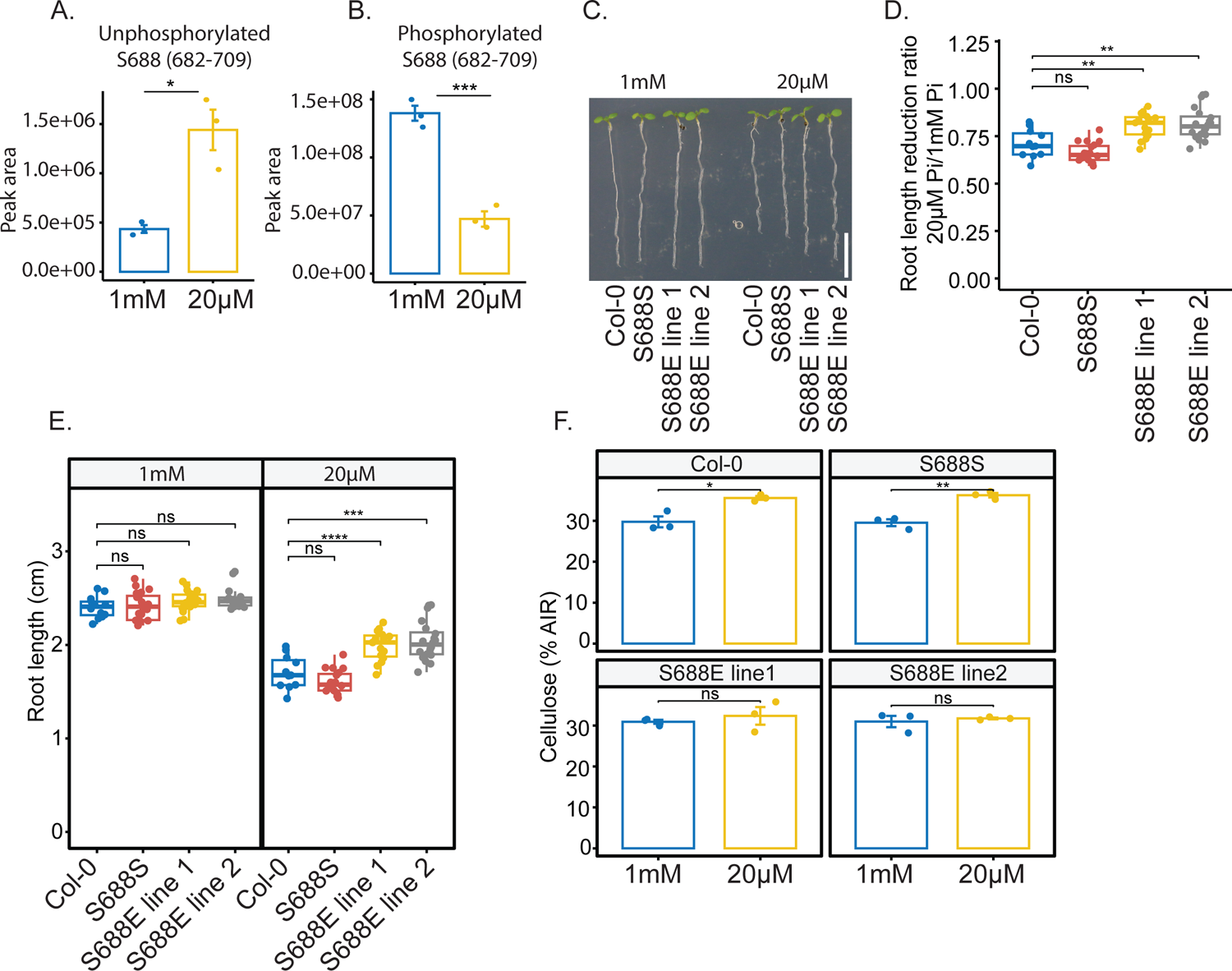
CESA1 phosphorylation is modified to regulate root growth under Pi deficient conditions. For A & B, roots were grown in a hydroponics system for 14 days in sufficient Pi (1mM) and transferred to Pi sufficient (1mM) or Pi deficient (20µM) media for 48h before experiments. **A** & **B**. Summed peak area of unphosphorylated (**A**) and phosphorylated (**B**) peptide containing S688 phosphosite. Data are presented as means ± SE, n = 3.**C.** Representative images of eight day-old seedlings grown on Pi sufficient and Pi deficient media. Scale bar = 1cm **D.** Primary root length reduction ratio of plants grown under Pi deficient conditions as compared to plants grown under Pi sufficient conditions. **E.** Primary root length of eight day-old plants grown under Pi sufficient and deficient conditions. **F.** Cellulose quantification in the eight day-old roots of plants grown under Pi sufficient and Pi deficient conditions. S688S and S688E lines are in the *cesa1* (-/-) background. Data are presented as means ± SE, n = 3. Asterisks denote statistical significance (*, P < 0.05; **, P < 0.01; ***, P < 0.001 and; ****, P < 0.0001) according to Student’s t test. The graphics and statistical analysis were generated using the ggpubR package in R.

**Figure 6.**
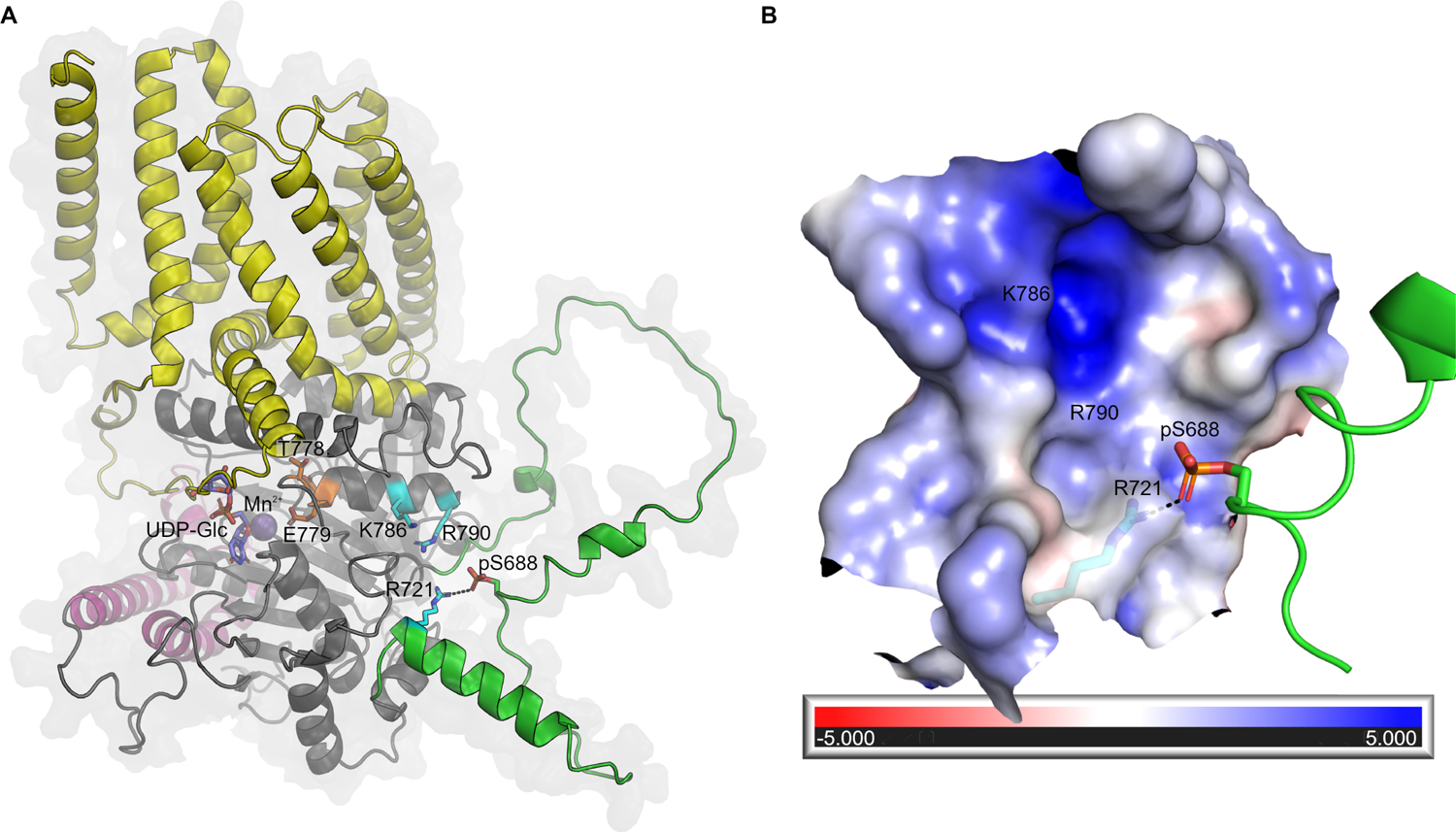
Structural basis for the effects of CESA phosphorylation. **A.** *At*CESA1 model showing the transmembrane domain (yellow), *Plant Conserved Region* (magenta) and CSR (green). Residues R721, K786 and R790 that contribute to the positive electrostatic potential in the pocket, the modelled phosphoserine (pS688) and TED motif (T778 and E779) are shown in cyan, green and orange, respectively. Black dashes indicate the possible interaction between pS688 and R721. The UDP-Glc (purple) and Mn^2+^ (sphere) ligands from the crystal structure of the *At*CESA3 catalytic domain (PDB 7CK2) are superimposed onto the *At*CESA1 model for comparison. **B.** The largely positive electrostatic potential of the pocket near the pS688 site is mapped onto the surface. The positions of residues R721, K786 and R790 are indicated.

### Phosphorylation of S688 in CESA1 might regulate the activity of the cellulose synthase complex via intramolecular conformational changes

To gain insights into a possible effect of the S688 phosphorylation of CESA1 function, we modelled the phosphoserine (pS688) and computed surface parameters. CESA1 S688 is located in the disordered part of the CSR (Q715-T649) modelled with low confidence which must be interpreted with caution. However, S688 is found immediately upstream in sequence from a reliably modelled α-helical segment (R716-W222) and modelling the pS688 indicate that its phosphate group could exist close to R721 (< 3 Å) in the vicinity of a cavity (∼ 12 Å) with positive electrostatic potential (formed largely by charged residues *e.g.* R721, K786 and R790) (Fig. 6AB). In addition, the flexibility of the partly disordered CSR region surrounding pS688 might allow it to interact with other charged residues in the nearby cavity *e.g.* K786 and R790, located on the same α-helix as the strictly conserved TED catalytic motif (Cruz *et al*., 2019, Pedersen et al., 2023). Thus, it is possible that the phosphorylation status of S688 may influence the catalytic activity of CESA1 via intramolecular conformational changes and/or affect the multimeric state of the CSC. Furthermore, we searched the FAT-PTM database (Functional Analysis Tools for Post-Translational Modifications, available at https://bioinformatics.cse.unr.edu/fat-ptm/), where we previously summarised phosphorylation sites in CESA proteins (Pedersen et al., 2023). Interestingly, we identified an equivalent documented phosphorylation site, pS671, in CESA3 (Wang *et al*., 2013). Additionally, equivalent charged residues R706, K771, and R775 were found to be present in CESA3, perhaps suggesting that this phosphorylation site might regulate the activity of CESA3 in a similar manner to CESA1.

## Discussion

Our study uncovered that Arabidopsis roots had higher cellulose content under Pi limited conditions (Fig. 1A). Usually, an increase in cellulose synthesis is associated with increased plant growth and cell expansion because new cell walls are constantly needed for cells to divide and expand. However, low Pi levels lead to a decrease in both cell division and cell elongation in primary roots (Abel, 2017). As a result, the increased cellulose deposition during Pi starvation likely results in stronger cell walls with reduced flexibility, negatively impacting cell expansion and growth. The cell wall thickness also increased at the cell elongation zone, which is consistent with previous research showing that cell wall thickness increases in response to Pi deprivation (Muller *et al*., 2015). It is important to note that normalisation of cellulose quantification with alcohol insoluble residue (AIR) may introduce artifacts, as reduced accumulation of another substance under Pi-starved conditions can falsely appear as increased cellulose content. However, our conclusions are supported by two other lines of evidence, including an increase in CSC speed under Pi starvation and a lack of cellulose increase in CESA1 phosphomimic lines.

An increase in cellulose content under Pi-deficient conditions is likely due to an increase in CSC activity at the plasma membrane. An increase in CSC activity has been documented to alter cellulose content (Sánchez-Rodríguez et al., 2017, Chen et al., 2010). One regulatory aspect of CSC speed involves changes in the phosphorylation of CESA proteins (Speicher *et al*., 2018a). Under low Pi conditions, the phosphorylation status of CESA1 proteins at the S688 site is reduced (Fig. 5AB). This is consistent with previous research showing that reducing the phosphorylation at the T157 site of CESA1 increases CSC speed and cellulose content (Sanchez-Rodriguez et al., 2017). It is, therefore, probable that the S688 site plays a similar role in CESA1 activity. In our study, the phosphonull S688A mutation was unable to rescue the *cesa1* knockout mutants. A previous study reported that the S688A mutation of CESA1 partially complements a conditional *cesa1* mutant, *rsw1-1*, leading to a decrease in CESA1 activity and cellulose content (Chen et al., 2010). However, the *rsw1-1* mutant is not a *cesa1* null mutant, making it difficult to interpret the outcome of the S688A version of CESA1 when another conditionally functional version of CESA1 is also present. The same study also found that the S688E version of CESA1 fully complements the *rsw1-1* (Chen et al., 2010). Similarly, we were able to effectively complement the *cesa1* knockout mutants with a phosphomimetic version of S688, suggesting that some level of phosphorylation is necessary for the CESA1 function. In addition, we discovered that the S688E version of CESA1 is less responsive to low Pi in terms of primary root growth reduction and cellulose synthesis (Fig. 5CDE). These findings imply that phosphorylation at the S688 site likely plays a role in regulating CESA1 activity, which in turn might control primary root growth, under Pi deficiency. It is possible that this is the result of altered CESA complex integrity, given that S688 is found in the CSR, which has a putative role in the oligomerization of the CESA complex. In addition, although pS688 is located in a disordered region sub-optimal for structural interpretation, it is bordered by a structurally ordered region (high model confidence) and the intermolecular CSR-CSR interface, which could retain pS688 approximately near the positively electrostatic cavity. Therefore, it is also possible that pS688 could interact with K786 and/or R790 located on the same α-helix as the conserved TED catalytic motif. We speculate that such interactions could lead to intramolecular conformational changes near the CESA1 active site, altering its activity.

Our phosphoproteomics analysis revealed significant changes in phosphorylation status of proteins involved in protein trafficking, specifically the ADP-ribosylation factor (ARF) GTPase protein family (Fig. 4C). These proteins play a role in orchestrating intracellular protein trafficking along the secretory pathway, particularly from the endoplasmic reticulum (ER) and Golgi apparatus to the plasma membrane (Donaldson & Jackson, 2011). The intricate process of intracellular trafficking is essential in facilitating Pi transport signalling. Notably, PHOSPHATE FACILITATOR1 (PHF1) is a key regulator governing the secretion of phosphate transporters (PHTs) to the plasma membrane (González *et al*., 2005). Mutations in the *PHF1* gene is associated with the retention of PHT1;1 in the ER, leading to reduced accumulation of this transporter in the plasma membrane (González et al., 2005). Consequently, these molecular perturbations result in diminished levels of Pi in plants and a constitutively active Pi starvation response. Furthermore, protein trafficking may be essential for adjusting the abundance of Pi transporters at various organelles, thus regulating the distribution of phosphate across cell compartments. Additionally, PHOSPHATE1 (PHO1), a key phosphate exporter involved in xylem loading, predominantly localises to the Golgi apparatus (Arpat *et al*., 2012). It is hypothesised that PHO1 is only transiently present at the plasma membrane due to its potential toxicity to plants (Arpat et al., 2012). Hence, precise regulation of protein trafficking is necessary to maintain an appropriate quantity of Pi transporters at the plasma membrane, ensuring Pi homeostasis. The identification of post-translational regulations in protein trafficking processes within our dataset presents a valuable opportunity to gain further insights into the mechanisms governing the maintenance of phosphate homeostasis.

Kinases regulate changes in root growth in response to Pi deficiency, e.g. mutants of CBL-interacting protein kinases (CIPK) are more sensitive to Pi deficiency in terms of root elongation (Lu *et al*., 2020). While CESA proteins can be phosphorylated at multiple sites (Speicher et al., 2018b), the specific kinases that mediate this phosphorylation have not yet been identified. It is likely that protein kinases involved in Pi starvation signalling also regulate CESA phosphorylation to control primary root growth under Pi-deficient conditions. CESA1 phosphorylation is regulated by the brassinosteroid (BR) pathway (Sanchez-Rodriguez et al., 2017). Here, the BIN2 protein, which is a key inhibitor of the BR pathway, impacts crystalline cellulose levels by directly phosphorylating CESA1 (Sanchez-Rodriguez et al., 2017). By contrast, the BES1 and BZR1 proteins, which are important transcription factors in the BR pathway, increase cellulose synthesis by binding to *CESA* promoters (Xie *et al*., 2011). In addition to directly regulating cellulose synthesis, BR also affects the rearrangement of microtubules, which are necessary for the proper direction of CSCs during cellulose synthesis (Catterou *et al*., 2001). The CELLULOSE SYNTHASE-INTERACTIVE PROTEIN 1, CELLULOSE SYNTHASE MICROTUBULE UNCOUPLING, and COMPANION OF CELLULOSE SYNTHASE proteins are involved in the connection of CSCs to microtubules (Bringmann *et al*., 2012, Liu *et al*., 2016, Endler *et al*., 2015). Interestingly, the BR signalling pathway also regulates primary root growth under Pi deficiency (Singh *et al*., 2018). The constitutive mutants of BRASSINAZOLE RESISTANT 1 (BZR1), a key regulator of BR-mediated regulation of gene expression (Yin *et al*., 2002), does not show root growth reduction under low Pi (Singh et al., 2018). This is likely due to the BZR1-mediated regulation of LPR1 but may also involve cellulose synthesis regulation under low Pi (Singh et al., 2018). Another study found that CELLULOSE SYNTHASE-LIKE B5 (CSLB5) is differently regulated in seedlings experiencing Pi deficiency. Further investigation revealed that plants with a mutation in the *CSLB5* gene had shorter root hairs than wild-type plants when grown in low Pi conditions, suggesting that CSLB5 plays a role in the plant’s response to low Pi levels (Lin et al., 2011). While the CSLB5 protein is part of the cellulose synthase superfamily, the CSLBs are currently not associated with any specific functions.

Apart from cellulose, other cell wall-related processes may be regulated in response to Pi deficiency. The transcriptomic comparison of three Pi response mutants (*lpr1*, *lpr2*, and *pdr2*) revealed significant changes in the expression of genes related to the cell wall in response to low Pi levels (Hoehenwarter *et al*., 2016). These changes occurred in four main areas: pectin modification, cell wall relaxation, hemicellulose/cellulose modification, and carbohydrate hydrolytic enzymes (Hoehenwarter et al., 2016). Data from several transcriptomic experiments show that low Pi levels lead to changes in the transcript abundances of numerous cell wall-related genes in Arabidopsis plants (Lin *et al*., 2011, Wu *et al*., 2003, Misson *et al*., 2005, Wege *et al*., 2016). Furthermore, low levels of Pi in plants result in an accumulation of lignin and callose in the cell walls of roots in an LPR1 dependent manner. Lignin is a structural component of the cell wall that helps to strengthen it by attaching to cellulose, but an excess of lignin can cause the cell wall to become less flexible. Pi deficiency can also cause an increase in the concentration of unesterified pectins in the cell walls of roots, particularly in the elongation zone and the root apical meristem (Hoehenwarter et al., 2016). These pectins can bind to Fe^3+^, Al^3+^, and Ca^2+^, and can form complex structures called “egg-boxes” through Ca^2+^-pectate crosslinking (Grant *et al*., 1973). The formation of egg-boxes can lead to the stiffening of the cell wall and reduced growth (Chebli & Geitmann, 2017).

Moreover, the cell wall serves as the primary interface between the cell and its external environment, making it the initial recipient of any alterations in nutrient availability. Local Pi sensing, which is primarily regulated by LPR proteins, prominently takes place within the plant cell walls (Muller et al., 2015). Remarkably, the efficacy of local Pi sensing relies on the presence of external iron (Fe) (Muller et al., 2015). LPR proteins utilize Pi-dependent cues to monitor subtle disparities in Fe availability in the apoplast (Naumann *et al*., 2022a).

Furthermore, plants employ modifications in the cell wall structure to release P that is bound to the cell wall under conditions of Pi deficiency (Zhu *et al*., 2014). It is probable that the presence of carboxylate groups (-COO-) in homogalacturonans (a type of pectin) enables the sequestration of Fe ions through the formation of -COO-Fe linkages. Under low Pi conditions, the increased abundance of pectins provides an elevated number of carboxylate groups, leading to enhanced binding of Fe and subsequent liberation of trapped phosphate from FePO_4_ complexes. The release of Pi from the cell wall is expected to exert a notable influence on the overall signalling response to Pi starvation. For example, in rice (*Oryza sativa*), mutants with cell wall synthesis defects show a constitutive Pi starvation response, and increased Pi transport was observed even when grown in high Pi conditions (Jin *et al*., 2015). In conclusion, our findings, combined with previous observations, emphasise the significance of cell wall dynamics in orchestrating the plant’s response to limited Pi availability.

## Material and Methods

### Plant material and growth conditions

The *prc1-1* mutant used in these experiments was previously described (Fagard et al., 2000). The *cesa1* T-DNA insertion mutant line (SAIL_278_E08) was obtained from the Arabidopsis biological resource centre (ABRC, https://abrc.osu.edu/). The homozygosity of the *cesa1* mutation was confirmed by PCR using the following forward and reverse primers: 5′-CAGAAGTGACTCCAATGCTCC-3′ and 5′-TGGTTGGTGGAATGAGAAGAG-3′.

A vertical agar plate method was used to conduct experiments with varied Pi conditions. The standard media contained half-strength MS (Murashige and Skoog, 1962) Basal Salt mixture (M407, Phytotech), 0.5% (w/v) MES (Sigma-Aldrich), 0.5% (w/v) Suc (Sigma-Aldrich), 10.3 mM NH_4_NO_3_, 9.4 mM KNO_3_, 0.05% (w/v), and 0.8% (w/v) Difco granulated agar (lot 6173985). For Pi sufficient conditions, standard media was supplemented with 1mM KH_2_PO_4_; for Pi deficient conditions, the standard media was supplemented with 20µM KH_2_PO_4_ and 980µM KCl. Arabidopsis seeds were surface sterilized in chlorine gas for three hours, resuspended in 0.1% (w/v) agarose, stratified at 4°C for two days, and then seeded in a single row on 12-cm-square plates that were sealed with 3M Micropore tape. Plates were arranged vertically into racks in the growth chamber, which had a 16-h light/ 8h-dark cycle, 120 E m2 s1 of light intensity, 23 °C (day)/19 °C (night), and 60% humidity.

### Cellulose quantification

Arabidopsis seedlings were grown on Pi sufficient (1mM) and Pi deficient (20µM) conditions for ten days. The roots were separated from the shoots, and crystalline cellulose was measured as previously described (Sanchez-Rodriguez *et al*., 2012).

### Electron microscopy

Roots were grown on Pi sufficient (1mM) and Pi deficient (20µM) for eight days, and the imaging was performed in the root elongation zones as described below. Eight-day-old Arabidopsis seedlings were fixed in 2.5% glutaraldehyde in phosphate-buffered saline (PBS) overnight at 4°C. The plants were washed 3 times in PBS, once in double distilled H2O (ddH2O) and then post-fixed in 1% osmium tetroxide for two hours. Following post-fixation, the samples were washed three times in ddH2O followed by gradual dehydration in 10% increments of acetone for one hour each to 100% acetone, where fresh acetone was exchanged three times. Following dehydration, the samples were infiltrated with Spurr’s resin (Spurr, 1969) in 25% increments for 8-12 h to 100% resin, which was again exchanged three times before embedding and polymerization. Cross sections of the embedded roots were then thin-sectioned using a Leica Ultracut 7 ultramicrotome (Leica Microsystems), collected on copper grids and stained with 1% aqueous uranyl acetate and Reynold’s lead citrate. The sections were imaged on a Tecnai Spirit transmission electron microscope equipped with an Eagle CCD camera. The cell wall thickness was measured using FIJI (Schindelin *et al*., 2012).

### Live cell imaging and data processing

Transgenic lines expressing *pCESA6*::*YFP-CESA6* have been previously described (Paredez et al., 2006). Seedlings for image analysis were grown for eight days in Pi sufficient or Pi deficient conditions. Roots were mounted in water under a pad of 0.8% agarose (Bioline). Imaging was performed using a CSU-W1 spinning disk head (Yokogawa) mounted to an inverted Nikon Ti-E microscope equipped with a 100× TIRF oil-immersion objective (NA = 1.49) and a deep-cooled iXon Ultra 888 EM-CCD camera (Andor Technology). A timelapse consisting of 800-ms exposure every 10 s for 5 minutes was recorded. Fiji (NIH) plug-ins “stackreg,” and “bleach correction” with default settings were used for drift and bleaching corrections respectively. “kymograph evaluation” plug-in of the free tracking software FIESTA was used for CSC speed analysis (Ruhnow *et al*., 2011). More than 100 kymographs were analyzed, and the speed of CSCs were determined by estimating the slopes of their trajectories with a straight line.

To determine the density of CSCs at the plasma membrane, areas of interest without significant Golgi signals were selected using the free-hand selection tool. CSCs were automatically identified on 8-bit images using the Find Maxima tool in Fiji (Schindelin et al., 2012), using the same noise threshold for all images. The density of CSCs in each area of interest was calculated by dividing the number of particles by the area of the region.

### Mass spectrometry analysis

Plants were grown in sufficient Pi media (1mM) for high root biomass production as described previously (Hetu *et al*., 2005). After 14 days, plants were treated with Pi sufficient or Pi media for 48 hours, and roots were harvested and snap-frozen in liquid nitrogen for subsequent protein extraction. Roots were ground in a mortal pestle, and powder was resuspended in protein extraction buffer (50mM MOPS, 2mM EDTA and 2mM EGTA) containing one tablet of cOmplete EDTA-free protease inhibitor cocktail (Sigma) per 50 mL of buffer and one tablet of phosSTOP (Sigma) per 20 mL of buffer. After 10min centrifugation at 10,000g to remove the debris, the supernatant was centrifuged at 80,000g for one hour to retrieve the membrane fraction in the pellet. The pellet was resuspended in the protein denaturation buffer (6M Urea and 100mM ammonium bicarbonate), and protein quantification was performed with Bicinchoninic Acid (BCA, Thermofisher) assay kit according to the manufacturer’s instructions.

250µg of proteins were reduced with 10mM DTT (Dithiothreitol) at 60°C for one hour and acylated with 15mM iodoacetamide (IAA) for 45min at room temperature. Urea concentrations were diluted to 1M with 100mM ammonium bicarbonate, and protein digestion was performed with ten µg of Trypsin per mg of proteins samples at 37°C overnight. Samples were acidified with formic acid and centrifuged at 10,000g for 10min at 4°C. The supernatant was desalted with SepPak C18 cartridges (Waters) according to manufacturer instructions. The desalted peptides underwent shotgun proteomics for label-free protein quantification and were also further processed using a titanium dioxide-based phosphopeptide enrichment method as described in (Sanes, 2019). LC-MS/MS was used to analyze the peptide samples. The LC system, Ultimate 3000 RSLC (Thermofisher Scientific, San Jose, CA, USA) was set up with an Acclaim Pepmap RSLC analytical column (C18, 100 Å, 75 μm × 50 cm, Thermo Fisher Scientific, San Jose, CA, USA) and Acclaim Pepmap nano-trap column (75 μm× 2 cm, C18, 100 Å) and controlled at 50°C. Solvent A was 0.1% v/v formic acid and 5% v/v dimethyl sulfoxide (DMSO) in water and solvent B was 0.1% v/v formic acid and 5% DMSO in ACN. The trap column was loaded with tryptic peptide at an isocratic flow 3% ACN containing 0.05% TFA at 6 µl/min for 6 min, followed by the switch of the trap column as parallel to the analytical column. The gradient settings for the LC runs, at a flow rate 300 nl/min, were as follows: solvent B 3% to 23% in 89 min, 23% to 40% in 10 min, 40% to 80% in 5 min, maintained at 80% for 5 min before dropping to 3% in 2 min and equilibration at 3 % solvent B for 7 min. A Fusion Lumos Orbitrap mass spectrometer (Thermo Fisher Scientific, San Jose, CA, USA) was employed at positive mode to execute the MS experiments with settings of spray voltages, S-lens RF, and capillary temperature level of 1.9 kV, 30 %, 275 °C, respectively. The mass spectrometry data was acquired with a 3-s cycle time for one full scan MS spectra and as many data dependent higher-energy C-trap dissociation (HCD)-MS/MS spectra as possible. Full scan MS spectra features ions at m/z of 400-1500, a maximum ion trapping time of 50 msec, an auto gain control target value of 4e5, and a resolution of 120,000 at m/z 200. An m/z isolation window of 1.6, an auto gain control target value of 5e4, a 35 % normalized collision energy, a first mass at m/z of 110, a maximum ion trapping time of 60 msec, and a resolution of 15,000 at m/z 200 were used to perform data dependent HCD-MS/MS of precursor ions (charge states from 2 to 5). Dynamic exclusion for 30 s was enabled. The results were analyzed using the Arabidopsis database TAIR10 and Protein Discoverer (version 2.4, Thermo Fisher Scientific) with Sequest HT. The ptmRS node was used in the analysis process. The mass tolerances were set at 10 ppm for precursor and 0.05 Da for fragment Mass Tolerance. The analysis allowed for up to two missed cleavages. Cysteine was fixed as a carbamidomethylation modification, while N-terminal protein acetylation and methionine oxidation were variable modifications. For phospho-peptides, additional parameters were set for variable modifications of phosphorylation on serine, threonine, and tyrosine. Label-free quantitation of the S688 containing peptide was performed using the Skyline software package (MacLean *et al*., 2010). Manual integration of the data was performed for peak integration (MacLean et al., 2010, Schilling *et al*., 2012). Peak areas were summed for three isotopic peaks (M, M + 1, M + 2) for each peptide to serve as the peptide’s quantitative measure. The mass spectrometry proteomics data have been deposited to the ProteomeXchange Consortium via the PRIDE (Perez-Riverol *et al*., 2021) partner repository with the dataset identifier PXD045341.

### Site specific mutagenesis and complementation

The wild-type sequence of *CESA1* was obtained from Arabidopsis cDNA through PCR amplification and cloned into the D-Topo entry vector from Invitrogen. Site-directed mutagenesis at the *CESA1* sequence was performed using the infusion cloning kit from Takara Bio, following the manufacturer’s instructions. Inverse PCR was used to create the S688A mutation with the following forward and reverse primers: 5′-AATGCTCCACTTTTCAATATGGAGGACATCGATGAGGG-3′ and 5′-TATTGAAAAGTGGAGCATTGGCGTCACTTCTGTTGATG-3′. The S688E mutation was created using the following forward and reverse primers: 5′-AATGCTCCACTTTTCAATATGGAGGACATCGATGAGGG-3′ and 5′-TATTGAAAAGTGGAGCATTTTCGTCACTTCTGTTGATG-3′. The entry vector was then transferred into a modified pMDC32 destination vector that contained a *CESA1* promoter. The 35S promoter in the pMDC32 vector was replaced with the *CESA1* promoter using the following method: the pMDC32 vector was linearised using the HindIII and KpnI restriction enzymes at the sites surrounding the 35S promoter. The *CESA1* promoter was amplified using the forward primer 5′-GGCCAGTGCCAAGCTGATTCTCATACGCCCGTCGA-3′ and reverse primer 5′-TCGAGGGGGGGCCCGCGCAGCCACCGACACACA-3′, which included a 15bp overlap at the extremities of the linearised pMDC32 vector. The amplified promoter and linearised pMDC32 vector were ligated using the infusion cloning enzyme from Takara Bio, following the manufacturer’s instructions. The ligation reaction was transformed into Stellar competent cells, and colonies were screened for correct pCesA1 insertion.

The *cesa1* homozygous knockout mutants are male gametophytic lethal; hence heterozygous plants were transformed with the chosen expression vectors with the floral dip method. For the selection of T1 plants, we utilized hygromycin as a selection marker for the transformed vector and performed PCR genotyping to identify heterozygous *cesa1* mutants. The selected plants were then self-fertilized to produce the next generation of seeds. T2 plants were selected for both homozygosity of the *cesa1* mutation by PCR genotyping and the presence of the transformed T-DNA by hygromycin selection in the next generation.

### Quantitative RT-PCR

The plant material was immediately frozen in liquid nitrogen and ground using the Invitrogen tissue lyser. Approximately 100 to 150 mg of powder was used to extract total RNA using the RNeasy Plant Mini Kit from QIAGEN. On-column DNase treatment was performed using the RNase-free DNase kit from Qiagen. 500 ng of RNA per sample and SuperScript III reverse transcriptase from Invitrogen were used for reverse transcription. Quantitative PCR was performed using SYBR select master mix from Invitrogen and primers described in the table S3.

### Protein modelling

The CESA1 model was retrieved from the AlphaFold (AF) Protein Structure Database (https://alphafold.ebi.ac.uk/files/AF-W8PUJ0-F1-model_v4.pdb). Parts of the CSR are modelled with low confidence scores (predicted local distance difference test, pLDDT < 70), indicating disorder. The position of the phosphoserine site S688 (albeit modelled with low confidence) is close in sequence to an α-helical segment (R716-E735, proximal to the catalytic domain) modelled with confident accuracy (pLDDT > 70). Phosphorylation was modelled in COOT v.0.9.8.1 (Emsley *et al*., 2010), choosing a rotamer with favourable geometry. Surface electrostatic potential was calculated with PDB2PQR (for partial charges and atomic radii) and APBS (Jurrus *et al*., 2018) (Adaptive Poisson-Boltzmann Solver) using 0.58 grid spacing.

## Acknowledgements

We would like to extend our gratitude to Joshua Heazlewood for his invaluable advice on proteomics. GAK was funded by the Swiss National Science Foundation Fellowship (P2LAP3-168408) and the ARC DECRA Fellowship (DE210101200). KEHF was funded by a Novo Nordisk Foundation Industrial Biotechnology and Environmental Biotechnology Postdoctoral grant (NNF21OC0071799) and a Villum Foundation Experiment grant (VIL50427). SP was funded by a Villum, two Novo Nordisk, and Danish National Research Foundation grants (25915, 19OC0056076, 20OC0060564, DNRF155, respectively). Furthermore, we acknowledge Mass Spectrometry and Proteomics facility at Bio21 Institute for their support in the processing of proteomics data. We also extend our appreciation to Julian Ratcliffe and the Bioimaging Platform of La Trobe University for their contributions to TEM imaging.

---

**Figure S1.** Primary root length measurement of segregating phosphomimic CESA1 (S688E) plants Root length of 8 days old seedlings grown on Pi sufficient (1mM) and Pi deficient (20µM) media. *cesa1* (+/-) seedlings were transformed with phosphomimic CESA1 (S688E) under the endogenous promoter. T1 plants were selected with hygromycin as a selection marker, and seeds from these plants were used for this experiment. ns =not significant according to Student’s t test. The graphics and statistical analysis were generated using the ggpubR package in R.

